# Integration of HLA-DR linkage disequilibrium to MHC class II predictions

**DOI:** 10.1101/2023.05.24.542040

**Authors:** Mette Bøge Pedersen, Sarah Rosenberg Asmussen, Freja Malou Sarfelt, Astrid Brix Saksager, Peter Wad Sackett, Morten Nielsen, Carolina Barra

## Abstract

Insights into peptide binding to HLA class II molecules is essential when studying the biological mechanisms behind cellular immunity, autoimmune diseases, and the development of immunotherapies and peptide vaccines. Currently, most of the publicly available data used to train state-of-the-art binding prediction methods for HLA-DR only includes DRB1 information. The role of the paralogue alleles, HLA-DRB3/4/5, and their strong linkage disequilibrium to DRB1 is often omitted when typing HLA-II alleles. This leads to ambiguities when making disease associations and interpreting HLA-restricted immune data. To resolve this issue, we present HLAAssoc-1.0, a method to infer HLA-DRB3/4/5 alleles by linkage disequilibrium to HLA-DRB1. We illustrate the usage of the tool and the importance of the integration of HLA-DRB3/4/5 alleles in the data analysis in different case studies including the interpretation of immunopetidomics data. Additionally, we infer allele information for the data used for training of NetMHCIIpan lacking HLA-DRB3/4/5 allele information and demonstrate that the retrained method achieved improved performance. In all cases, inferring HLA-DRB3/4/5 allele presence in non-fully typed HLA-II assays resulted in improved allele and motif deconvolutions. HLAAssoc-1.0 is available at https://services.healthtech.dtu.dk/service.php?HLAAssoc-1.0.

## Introduction

The Major Histocompatibility Antigen (MHC) class II molecules function as a cell surface receptor that presents peptide antigens on antigen presenting cells (APC) to circulating T cells. MHC-II is responsible for binding and exposing peptides derived from endocytosed antigens(1). Depending on the immunogenicity of the presented peptide, an immune response can be initiated as the CD4 T lymphocytes recognize and bind to the peptide-MHC complex(2). This will lead to the activation of other immune cells such as macrophages, B lymphocytes, and CD8 T lymphocytes, which will fight the infection(1).

The MHC II molecule consists of a noncovalent complex of two chains i.e., the α-chain and β-chain forming an open peptide binding groove allowing binding of peptides of variable length, typically of 12-25 amino acids(3,4).The binding pockets of the MHC II binding groove typically interact with the side chains of residues 1, 4, 6, and 9 of a 9-mer core region of the bound peptide. The interaction with the peptide core region is strongly determinant for the binding affinity and specificity of the MHC II molecule(5,6).

MHC-II or Human Leukocyte Antigen (HLA) in humans, is polygenic, meaning the chromosome 6 has multiple loci containing genes which encode to different HLA molecules(1,7). The three main loci encoding MHC II are HLA-DR, HLA-DP, and HLA-DQ(8). The HLA-DR loci includes HLA-DRA1, encoding the α-chain, while the β-chain is encoded by the principal HLA-DRB1 gene, and possibly one of the three paralogues; HLA-DRB3, HLA-DRB4 and HLA-DRB5(1,9). The polygenicity of the MHC molecule results in most humans having both HLA-DRB1 and -DRB3/4/5 genes encoding the β-chain, though in some cases the gene occupying the HLA-DRB3/4/5 locus is noncoding.

The HLA is one of the most polymorphic genes known. There are currently more than 4,300 known HLA-DRB alleles in the IPD-IMGT/HLA database(10). HLA polymorphism allows every individual to possess MHC molecules with a unique peptide binding repertoire which increases the population diversity of the immune system and the ability to recognize and fight a wider range of exogenous hazards.

Since the HLA-DRB1 gene is more abundantly expressed than the HLA-DRB3/4/5 genes(9), the role of the paralogous DRB3/4/5 alleles is highly understudied compared to DRB1. Additionally, the loci for the two genes encoding the β-chain (HLA-DRB1 and HLA-DRB3/4/5) are located close to each other, which results in a strong linkage disequilibrium (LD)(11–13). Because of these matters, earlier studies which have found HLA-DRB1 alleles in close relation to certain diseases, might have omitted the possibility of the HLA-DRB3/4/5 being the actual cause of effect(14). Recent studies suggest that the secondary HLA-DRB alleles play a significant role in antigen-presentation with different peptide binding preferences compared to HLA-DRB1 molecules(13,15,16). To understand the antigen-presentation and immune response, identifying the binding preferences and immunopeptidome contribution of secondary HLA-DRB alleles is therefore of crucial importance.

Several autoimmune diseases (AIDs) have shown to be associated with specific HLA-DQ and HLA-DR alleles, increasing or decreasing the risk of disease development(17). For example, previous studies have identified HLA-DRB haplotypes that act as genetic risk factors for developing multiple sclerosis (MS) and type-1 diabetes(18–21). Both HLA-DRB1*15:01 and HLA-DRB5*01:01 were highly associated with MS, but it remained unclear which allele played the most important role(22). For a long time is was assumed that HLA-DRB1*15:01 was the most important allele, as it has a higher protein abundance, however recent evidence suggests that the DRB5*01:01 also plays an important role(23).This is inline with the fact that the two DR15 allomorphs have similarities in their peptide binding motifs. Furthermore, MS and Sjögren’s Syndrome share several common susceptibility genes, including HLA-DRB1 and HLA-DRB5(24). In relation to type-1 diabetes, T cell responses against a DRB4*01:01 restricted preproinsulin peptide have been observed(25). By contrast, susceptibility genes for rheumatoid arthritis seem to be primarily associated with HLA-DRB1, but there is a protective factor associated with HLA-DRB5(26). In addition, several studies have shown that it is possible to stratify patients based on the expression level of the different HLA genes. For instance, in Vitiligo, HLA-DRB4*01:01 was observed to be negatively correlated with late onset of the disease(27). This evidence emphasizes the importance of including the secondary HLA-DRB alleles to improve immunotherapy treatments(1,28).

Here, we present the tool HLAAssoc-1.0 to resolve this need by inference of the association frequencies of HLA-DRB3/4/5 with HLA-DRB1 alleles. The tool is based on haplotype data from a wide population study, performed by the National Marrow Donor Program (NMDP), containing full HLA haplotype typing (including HLA-DRB3/4/5)(29). Currently, the majority of publicly available data used for the training of MHC-II predictors primarily contains HLA-DRB1 information, and therefore the scarce data on those alleles difficult the task of learning their binding preferences. Using HLAAssoc-1.0, we have integrated the HLA-DRB3/4/5 LD typing into NetMHCIIpan-4.1, a current state of the art MHC class-II predictor and showed a performance improvement on HLA-DRB345 predictions. Further, the influence of adding additional HLA-DRB3/4/5 information given by linkage disequilibrium was assessed and benchmarked against no-or random linkage with existing tools.

## Methods

### Haplotype frequency data

Linked HLA-DRB1∼HLA-DRB3/4/5 haplotypes and their frequencies in 26 different population categories were extracted from ‘Be The Match Registry’(29), a voluntary bone marrow donor dataset sequenced by the National Marrow Donor Program (NMDP). The data consisted of 4419 different HLA haplotypes obtained by DNA-typing of 6.59 million individuals divided in 5 broad ethnicity categories with a subset of 2.9 million individuals divided in 21 detailed population subgroups. The sample sizes for the different populations ranged from 347 for the Alaska Native / Aleut population up to 1,596,577 for the Caucasian population (Supp. Table 1).

Four alleles in the data contained the g suffix; DRB4*01:01g, DRB3*02:02g, DRB5*01:02g, DRB3*02:01g, meaning groups of non-disambiguated alleles. The alleles in a “g” group have identical nucleotide sequences for the exons making up the binding groove (exon 2 for MHC-II)(10). In particular, for HLA-DRB4 no alleles other than DRB4*01:01g were described, hence all DRB4 were denoted as DRB4*01:01. The g suffix was removed from the allele names neglecting the ambiguity.

The dataset also included haplotypes with the non-functional DRBX*NNNN annotation, corresponding to individuals expressing only the DRB1 allele.

### Defining a representative set of HLA-DRB molecules

A representative set of HLA-DRB molecules was defined based on protein sequence data from the IPD-IMGT/HLA database(10) (retrieved April 2022), which contained 3888 different HLA-DRB molecules. To select representative HLA-DRB alleles, first, only those that appeared in the data with full protein sequence were chosen. From these, molecules were further selected based on variations in their pseudo-sequences (defined in(30) as a set of 34 polymorphic residues in the HLA alpha and beta protein chains in structural proximity to the bound peptides) to highlight functional differences. In case of HLA-DR molecules sharing pseudo-sequence, the allele with the name having the lowest number in the second field was selected (e.g. DRB1*05:01). The pseudo-sequence is defined here as a set of selected residues of the HLA-DRB polypeptide chain that interact with the bound peptide. This reduction resulted in a set of 123 HLA-DRB alleles.

### Functional tree clustering based on predictions with NetMHCIIpan-4.0

NetMHCIIpan-4.0 was used to make HLA antigen presentation predictions for 100,000 random natural peptides for the representative set of HLA-DR molecules. Next, a functional distance matrix was constructed between each pair of HLA-DR using the MHCcluster approach(31). Here, in short, the distance is defined as one minus the correlation between predicted binding values of the union of the top 10% highest scoring peptides for a given pair of HLAs. 100 distance matrices were computed using bootstrap of which a final consensus distance matrix was generated and converted into an Unweighted Pair Group Method with Arithmetic mean (UPGMA) tree.

### Sequence logos

Sequence logos were created using Seq2Logo-2.0 with default settings(32). The sequence logos were created from the predicted 9-mer peptide binding cores of the top 1 % strongest binding peptides for each molecule.

### The HLAAssoc-1.0 method

The HLAAssoc-1.0 method takes two types of input. One type is ‘donor input’ containing donor IDs and HLA-DRB1 genotype information for each donor (Supp. Fig 1). Here, the secondary alleles are inferred to each of the HLA-DRB1 alleles in the donor. The output file provides the inferred HLA-DRB3/4/5 alleles and their frequencies for each HLA-DRB1 allele as provided in the input (Supp. Fig. 1). Alternatively, HLAAssoc-1.0 can be used to determine associated HLA-DRB3/4/5 alleles for a given input file with HLA-DRB1 population frequencies. In this instance, the frequencies of the associated HLA-DRB3/4/5 alleles are calculated to each HLA-DRB1 allele individually and then added when more than one association to the same secondary allele. Ultimately the relative frequencies of HLA-DRB3/4/5 alleles are estimated by multiplying the frequency of the HLA-DRB3/4/5 allele with the frequency of the associated HLA-DRB1 allele provided in the input file (Supp. Fig 2).

The output file can be filtered by either a predefined frequency threshold or a top number. By default, a threshold of 0 is used, meaning that all the associated HLA-DRB3/4/5 alleles will be included. If the user restricts the output to a top while inputting donor data, the server will only show the chosen top of the associated alleles meeting a frequency of at least 5%.

### HLA-DRB1 world population frequencies

HLA-DRB1 frequencies datasets were downloaded from Allele Frequency Net Database (AFND, http://www.allelefrequencies.net) on May 2022(33). Only populations with a sample size of n≥10,000 were included. The data was aggregated for the DRB1 alleles and the frequencies were made to sum to one.

The alleles HLA-DRB1*04:57, HLA-DRB1*14:54, and HLA-DRB1*15:19 were not found in the haplotype data, therefore allele linkage inference to DRB3/4/5 alleles is missing for those alleles.

### Retraining NetMHCIIpan with HLA-DR linkage disequilibrium inference

NetMHCIIpan-4.0 was originally trained covering 124 class II MHC molecules. 42 MS immunopeptidomics data set from the NetMHCIIpan-4.0 publication(34) were found to have incomplete secondary HLA-DR annotation. For those, HLA-DRB3/4/5 alleles were inferred using HLAAssoc-1.0. This resulted in an expanded allele information for 34 data sets. With extended allele associations in the allele lists provided by HLAAssoc-1.0 a NetMHCIIpan-4.1 was retrained following the same procedure as described for NetMHCIIpan-4.0(34). In addition, new updated 2747 DRB protein sequences were downloaded from IMGT/HLA database(10) and added to extend the predictions of NetMHCIIpan-4.1.

The updated method trained on the data with expanded HLA-DR secondary allele information is implemented as a tool termed NetMHCIIpan-4.1, and is available at https://services.healthtech.dtu.dk/services/NetMHCIIpan-4.1.

Likewise is the updated NetMHCIIpan-4.1 included in the MHCMotifDecon motif deconvolution tool(16) available at https://services.healthtech.dtu.dk/services/MHCMotifDecon-1.0/.

## Results

### HLA-DRB3/4/5 predicted binding motifs using NetMHCIIpan-4.0 showed divergent amino acid preferences when comparing to HLA-DRB1

As a first means to characterize the complementary function between HLA-DR molecules formed by HLA-DRB1 and HLA-DRB3/4/5 alleles observed earlier(16), a set of representative molecules were clustered using the MHCcluster approach(31). This method clusters MHC molecules based on their functional specificities of the 9-mer core of the peptide they bind, using a simple agglomerative hierarchical clustering (for details see materials and methods).

Most of the paralogue HLA-DRB3/4/5 representative molecules are grouped in divergent clusters with clearly distinct binding motifs when compared to HLA-DRB1 (Figure 1). The binding motif of HLA-DRB5, differently to HLA-DRB1, showed a strong preference for positively charged amino acids in position (P)9. HLA-DRB4 showed a strong preference for Glutamine (Q) in P4, and negatively charged amino acids in P7, altering the typical P6 anchor position for DRB1. HLA-DRB3 displayed more diversity on binding preferences showing both preferences for Asparagine (N) or Aspartate (D) in P4 (the last related to HLA-DRB1*03 preferences). Overall, these differential residue preferences suggest the evolutionary diversification of binding motifs.

**Figure 1.**
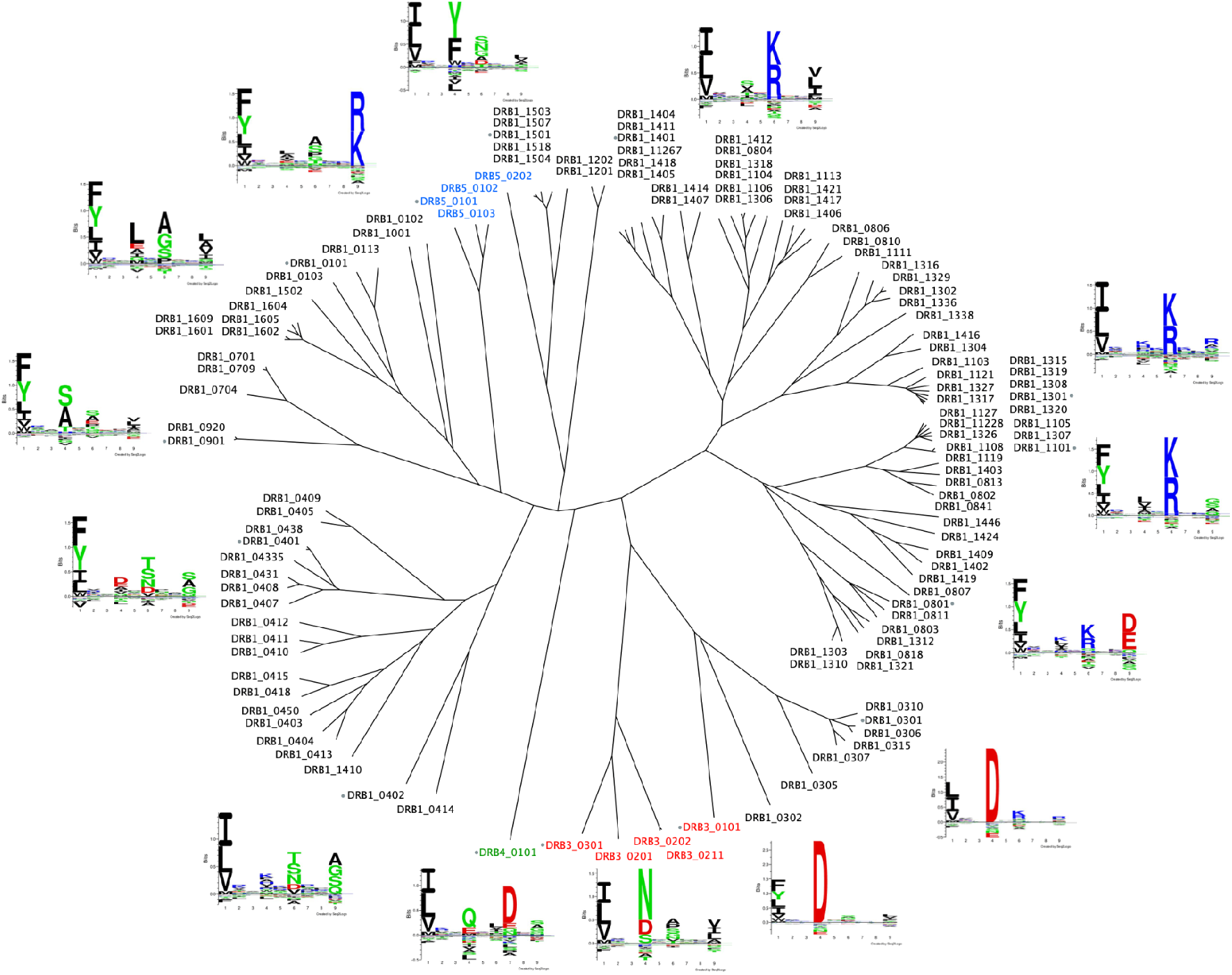
Functional tree clustering based on predictions with NetMHCIIpan-4.0. Peptide binding predictions for a representative set of HLA-DRB molecules were made by using NetMHCIIpan-4.0. The top 10% best binders of each allele were used to calculate distances between the alleles by using the MCHCluster method. 123 representative HLA-DRB molecules are visualized in an unrooted tree using SplitsTree4(35) and the sequence logos generated by Seq2Logo-2.0 are shown for alleles representing each cluster. The DRB1 alleles are colored black, DRB3 red, DRB4 green and DRB5 blue and a gray dot is added to the alleles being shown with a sequence logo.

### HLA-DRB3/4/5 can be inferred by linkage disequilibrium to HLA-DRB1

Given the importance of HLA-DRB3, 4, and 5 in complementing the functional space of HLA-DRB1, and the fact that this allele information is often unavailable, we developed HLAAssoc, a method that infers HLA-DRB3/4/5 alleles associated with HLA-DRB1 determined by linkage disequilibrium when full HLA-DR typing is absent. The method infers the associations using haplotype frequency data from the NMDP registry(29), which contains frequencies for 4,419 different HLA-DRB1∼HLA-DRB3,4,5 paired haplotypes typed in 26 different populations (refer to Methods section for more details). Based on this data, HLAAssoc calculates the frequency of HLA-DRB3/4/5 alleles associated with HLA-DRB1 genotypes in individual donors from specific populations. In short, the HLA-DRB3/4/5 allele(s) in disequilibrium with a given HLA-DRB1 in a given population are recorded and the expected allele frequency estimated from the corresponding haplotype frequency (for details refer to materials and methods). A world population was added to the list, by weighting the haplotype frequencies of each population using the sample size for the particular population, and then summing them all into one column of frequencies. The world population is used if no population is specified. In addition, HLAAssoc also allows to infer the HLA-DRB3/4/5 distribution in a population cohort with available HLA-DRB1 frequencies.

An implementation of the tool is available as a web service (Figure 2) at https://services.healthtech.dtu.dk/service.php?HLAAssoc-1.0, and can be downloaded on the web service page, to be implemented on bioinformatic pipelines.

**Figure 2.**
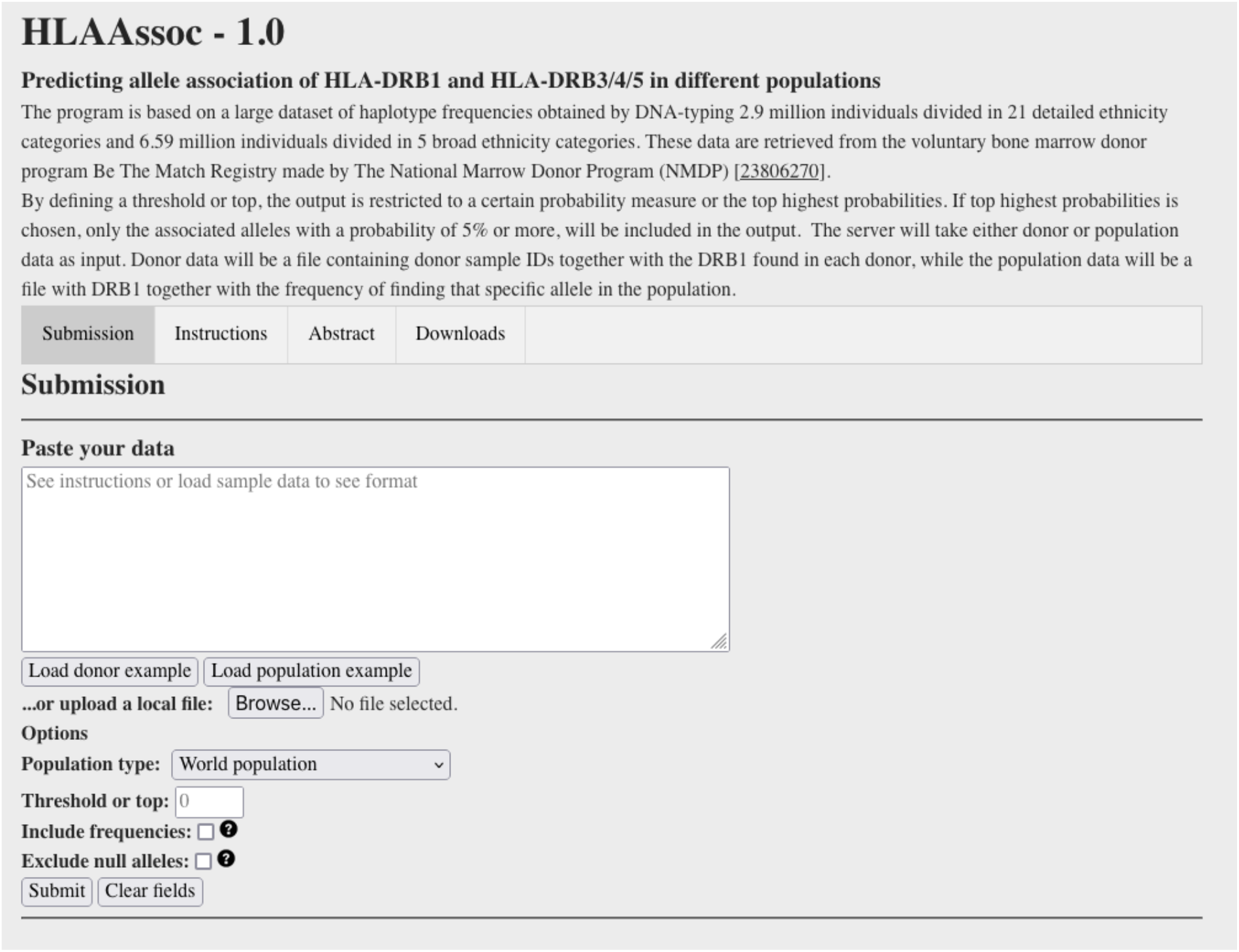
HLAAssoc-1.0 server interface. The HLAAssoc-1.0 method takes donor or population input modes to infer HLA-DRB3/4/5 associated to HLA-DRB1 in linkage disequilibrium. In population input mode, HLA-DRB1 frequencies should be provided.

The server takes two different types of input. One type is donor input containing donor IDs and DRB1 genotype information for each donor. The other is population input, which is a list of HLA-DRB1 alleles found in a population together with their frequencies. If the input is donor data, the output file will include the input data in the first three columns. The inferred alleles and their frequencies will be added as four additional columns; a column of inferred HLA-DRB3/4/5 or null alleles from each primary HLA-DRB1 allele, and a column of corresponding frequencies per HLA-DRB3/4/5 allele association. The associations are ordered by highest frequency. If the input uses population mode, e.g. all HLA-DRB1 frequencies in a population, the output will be a list of all the associated DRB3/4/5 to HLA-DRB1 with their corresponding frequencies (Ordered by highest frequency).

As an example of the use of the method, HLA-DRB3/4/5 frequencies were inferred based on world population HLA-DRB1 frequencies estimated from Allele Frequencies Net Database(36). Here, HLA-DRB1 frequencies for 33 different populations (all available populations with sample size≥10,000) were aggregated as described in methods (Supp. Table 1). The aggregated HLA-DRB1 frequencies were submitted to HLAAssoc-1.0 using “World population” and selecting the top 10 (Supp. Table 2).

In this analysis, HLA-DRB4*01:01 was found to be the most abundant secondary DRB allele with a frequency of nearly 30%, followed by HLA-DRB3*02:02 (21.7%), and the null allele (14%), denoted here as DRBX*NNNN (Fig. 3A). Summing up HLA-DRB3, HLA-DRB4 and HLA-DRB5 alleles per loci, we find that the most frequent is HLA-DRB3 which is observed in 40% of the world population (Fig. 3B), while HLA-DRB5 and the absence of a secondary DR beta allele (DRBX), have the lowest frequencies of around 14-15%.

**Figure 3.**
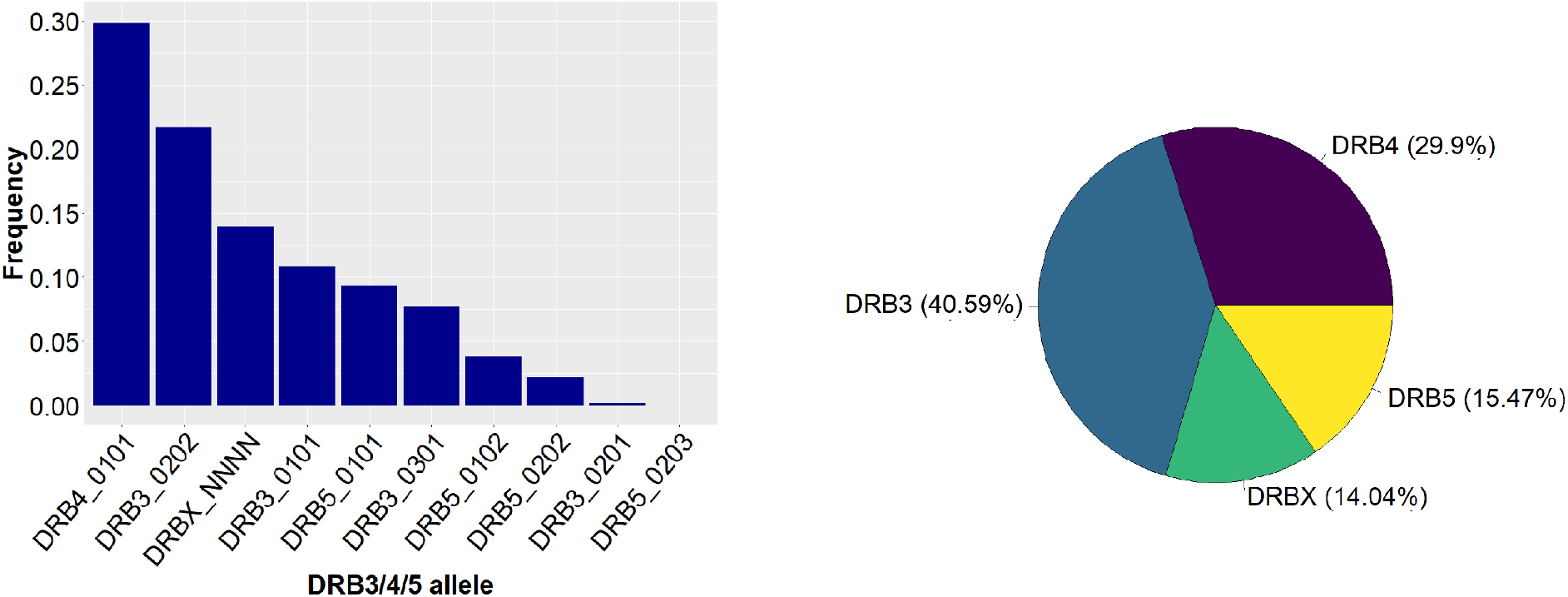
Distribution of top 10 DRB3/4/5 alleles in the world population. The distribution of HLA-DRB3/4/5 in the world population estimated from the aggregated HLA-DRB1 frequencies for all populations with sample size≥10,000 from allelefrequencies.net into HLAAssoc-1.0. **A**. Top 10 most frequent DRB3/4/5 alleles in the world population. **B**. Distribution of HLA-DRB3, -DRB4, -DRB5, and the non-functional allele -DRBX in the world population. *Note that the frequency for DRB4*01:01 might not be representative for this allele because of its ambiguity, instead the value denotes frequency of any DRB4 allele, hence denoted DRB4*01:01g in the original data.

### Retraining of NetMHCIIpan -Impacts on predictive power

Next, we assessed the impact of extending the HLA-DRB1 annotations with inferred HLA-DRB3/4/5 for the prediction of peptide HLA-DR presentation. Here, models for HLA class II antigen presentation were trained on the NetMHCIIpan-4.0 data set. 42 MS-immunopeptidomics data set from the NetMHCIIpan-4.0 training data were found to have missing secondary HLA-DR annotation. For these data sets HLA-DRB3/4/5 alleles were inferred using HLAAssoc-1.0 with a threshold of 0.5 (only associations with higher frequencies were included). This resulted in an expanded allele information for 34 data sets (Supplementary Table 3). Next, two models, one including the expanded allele information for these data sets (N) and one keeping the original information (O) were trained and evaluated using 5-fold cross-validation as described in the original NetMHCIIpan-4.0 publication, and performance estimates calculated in a per data set manner. Here, the data sets were split in two groups corresponding to data sets with expanded (Updated, or Up) HLA-DRB3/4/5 annotation and data sets without expanded allele information (Figure 4 and Supp. Fig. 4).

**Figure 4.**
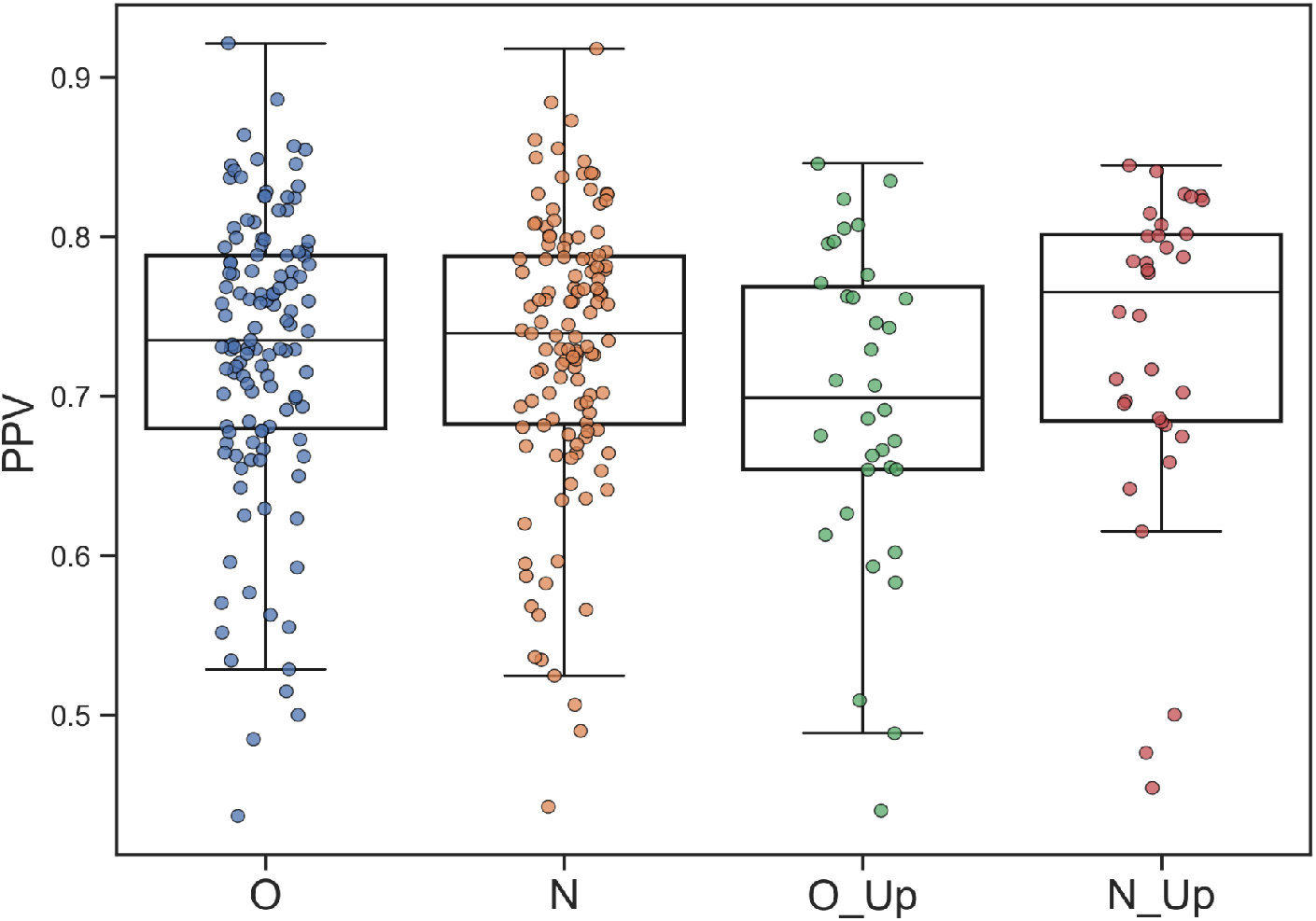
Cross-validated performance of models trained with original (O) and new (N) allele-list annotation. Data set is split into two subsets. One with entries without change of allele-list information and one with entries with expanded (labeled _Up) allele-list information. The gain in performance between model O and N is statistically significant for all three performance measures when comparing the UP data subset (p<0.009, binomial test excluding ties in all cases), and likewise borderline significant (p<0.1, binomial test excluding ties in all cases) for the remaining data subset. Positive predicted value (PPV). Here defined as the proportion of true positives within the top N predictions, where N is the total number of positives in the given data set.

These results demonstrate a consistent and highly statistically significant performance gain for the model trained on the extended allele information for the data sets with altered alleles (N_up versus O_up). In addition, the model trained with extended HLA-DR3/4/5 associations further showed a performance improvement for the data sets without altered allele information, though not statically significant (p<0.1, Figure 4 and Supp. Fig 4). These results underline the critical importance of including a complete allele annotation when analyzing HLA-DR immunopeptidomics data.

### Adding HLA-DRB3/4/5 annotations in MS immunopeptidomics deconvolution analysis rescued a significant number of peptides from the trash cluster

We next investigated how motif deconvolution of HLA-DR immunopeptidomics mass spectrometry (MS) was affected when excluding information related to HLA-DRB3/4/5 encoded HLA molecules. Here, we collected data from 40 fully HLA-DR typed donors from an earlier publication(37) (data available at https://services.healthtech.dtu.dk/suppl/immunology/Immunology2020_Attermann).

We cloistered HLA-DRB3/4/5 allele annotation, and used HLAAssoc-1.0, to infer HLA-DRB3/4/5 associations in linkage disequilibrium to HLA-DRB1 (Supp. Table 4). Using top 1 and world population for the association, HLAAsoc inferred the correct secondary DRB alleles with 90% (36/40) correct annotations as the inferred alleles are most often found in very strong linkage association. Most inferred alleles are based on linkage associations of 90% or higher. However, four cases resulted in an incorrect secondary HLA-DRB3/4/5 inference, all associated with DRB3*02:02. Three of the cases stems from an inference from DRB1*13:01. This allele is in linkage disequilibrium with two DRB3 alleles, DRB3*01:01 (58%) and DRB3*02:02 (42%), and in the three failed cases the less frequent allele was observed in the donor. The other case stems from DRB1*03:01. This allele is in linkage disequilibrium with the same two DRB3 alleles with frequencies of 83% and 17% respectively. Also here, the lower frequency allele was observed in the donor (Supp. Table 4).

Next, MHC motif deconvolution was performed for each of the 40 donors using MHCMotifDecon-1.0 (which builds on NetMHCIIpan-4.1 binding predictions) with the original full HLA-DR typing, with only DRB1, and with inferred HLA-DRB345 allele annotations for the complete immunopeptidome data set. Figure 5 gives the resulting motif deconvolution for two selected examples. Figure 5A displays an example where the inferred secondary DR alleles align with the observed (Donor11), and Figure 5B an example where an additional incorrect DRB3 was inferred (Donor15).

**Figure 5.**
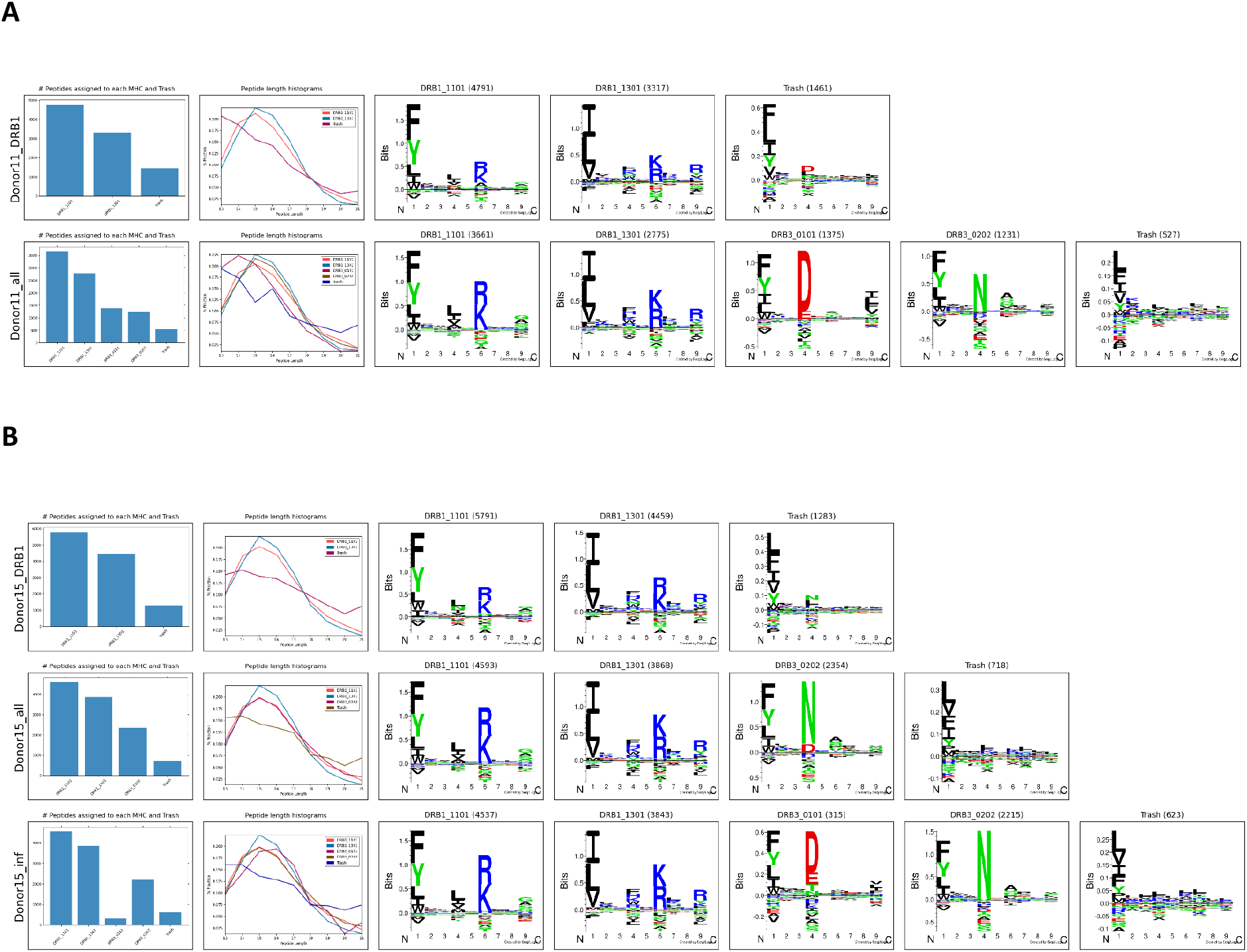
Motif deconvolution of MS dataset with and without HLA-DRB345 inference. Motif deconvolution was performed using MHCMotifDecon for no-and inferred/real allele associations. The barplots show the counts associated with the different alleles or the trash cluster (peptides with rank score higher than 20% for any of the present alleles). The length histograms show the distribution of the peptide length predicted for each of the alleles. ‘Donor_DRB1 ‘shows the deconvolution including only HLA-DRB1 alleles, ‘Donor_all’ shows the deconvolution including the full HLA-DR typing, and ‘Donor_inf’ shows the deconvolution based on the inferred secondary HLA-DRB3/4/5 (lower panel). **A**. Donor 11, and **B**. Donor 15.

Here, we observed that the motif deconvolution becomes more accurate and captures a higher proportion of the input immunopeptidome when adding the secondary HLA-DRB3/4/5 allele information (∼64% of the peptides are captured in the two examples when including only HLA-DRB1, and ∼82% when expanding to also include HLA-DRB3 in the two examples, using a binding threshold of 5% rank). Furthermore, we observe a very low proportion of peptides assigned to the wrongly inferred HLA-DRB3 (Fig. 5B) suggesting that incorrect inference can be corrected when combining the inferred predictions with motif deconvolution tools.

The increased proportion of captured HLA-DR ligands observed when including the paralogues HLA-DRB3/4/5 is to some extent trivially expected since we add the number of molecules to which we can predict binding. To address if this increase on HLA-DR restricted peptides is due to a true peptide-HLA association and not due to a spurious association using MHCMotifDeconvolution, we further quantify how the inclusion of secondary random HLA-DR alleles resulted in an improved motif deconvolution and compared to when using inferred associations identified by HLAAsoc. For each donor in the previous dataset, we performed MHC motif deconvolution adding the same number of random HLA-DRB3/4/5 alleles as the number of the assigned inferred alleles using HLAAssoc. MHC motif deconvolution was performed for each donor based on this allele list (Figure 6). The results showed that the motif deconvolution using the correct HLA-DRB3/4/5 alleles (Real or Inferred) captured more HLA-DR ligands for each donor and HLA-DR allele annotation (Figure 6).

**Figure 6.**
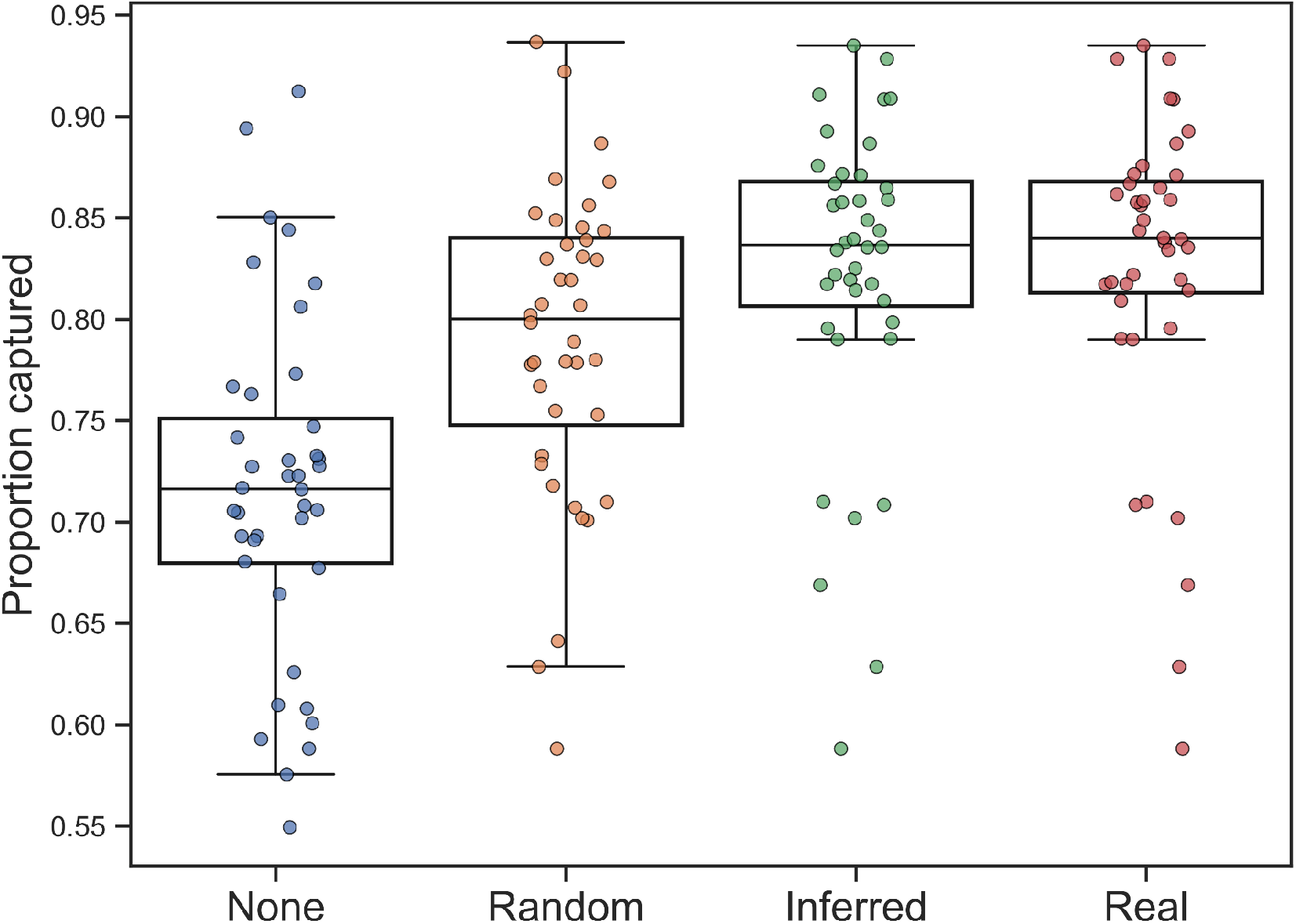
Proportion of captured peptides for each of the methods to annotation of secondary HLA-DR alleles. A captured peptide is defined as a peptide with a predicted percentile rank score of 5% or less to one or more of the HLA-DR alleles associated with a given Donor data set, and the reported performance is the proportion of such predicted binding out of the total pool of peptides for a given Donor. Each point in the plot corresponds to one donor. The different methods are “None”: No HLA-DRB345 alleles included, “Random” random HLA-DRB345 alleles associated, “Inferred”: HLA-DRB345 alleles inferred using HLAAssoc-1.0, and “Real”: The typed HLA-DRB345 alleles.

This analysis confirmed the findings that although the number of associated HLA-DR restricted peptides increased by adding more alleles that can bind the trash peptides, the inferred and real associations capture more. Overall, the proportion of captured HLA-DR ligands was increased when including the secondary DR alleles and the gain was found to be comparable when using either inferred or real DR allele associations, and significant (p<0.0001, binomial test) compared to random associations.

## Discussion

In this study, we have developed a method called HLAAssoc-1.0, for inferring secondary HLA-DRB3/4/5 alleles from HLA-DRB1 information based on haplotype frequency data that includes HLA-DRB1∼HLA-DRB3/4/5 paired typing in 26 different populations. The method utilizes linkage disequilibrium statistics to infer HLA-DRB3/4/5 alleles associated with HLA-DRB1 alleles. The dataset used for developing this model consists of 4,419 different haplotypes, allowing for the inference of secondary HLA-DR alleles across different populations.

Our findings align with previous work(16), demonstrating that secondary HLA-DRB alleles play a complementary role in antigen presentation, exhibiting different epitope preferences compared to HLA-DRB1. We observed that the secondary HLA-DRB3/4/5 molecules form distinct clusters, differing from HLA-DRB1 in functional-based clustering.

Here, we have demonstrated the accuracy of the tool in inferring the correct DRB3/4/5 alleles using a benchmark dataset. We applied the tool to expand the HLA-DR allele annotation for the training data of NetMHCIIpan, including information about HLA-DRB3/4/5 for datasets where this information was previously absent. This resulted in significant improvement in prediction accuracy for the expanded allele annotation datasets, strongly indicating the important contribution of secondary HLA-DR alleles in characterizing the immunopeptidome data used for training and evaluating NetMHCIIpan.

Building upon the improved version of NetMHCIIpan (version 4.1), we employed the MHCMotifDecon method developed for motif deconvolution on a large dataset of immunopeptidome data. We demonstrated how the inference of secondary HLA-DR alleles led to greatly enhanced deconvolution, reflected in an increased proportion of observed ligands being assigned with high likelihood to sample-specific HLA-DR molecules. This result further underscores the significant contribution of secondary HLA-DR molecules to the immunopeptidome.

However, it’s important to note that since HLAAssoc-1.0, in its current version, is constructed from data obtained in a single study, its performance may be susceptible to bias and a lack of data from certain populations. This means that certain associations of secondary HLA-DR alleles might be overlooked when using the tool. One example of such bias is related to DRB4 alleles, where ambiguity exists, and they are represented in the format DRB4*01:01g. Consequently, the HLAAssoc tool maps all DRB4 inferences to DRB4*01:01. We anticipate that the impact of this issue will diminish as more diverse cohort haplotype data becomes available and is integrated into the tool.

Currently, the HLAAssoc tool is limited to HLA-DRB3/4/5 allele inference. However, there is linkage disequilibrium between other HLA loci, including HLA-DR and HLA-DQ(38), as well as between HLA-A/B and HLA-DR, and within HLA class I(39). A natural future extension of the HLAAssoc tool would be to incorporate these associations as well.

In conclusion, we have developed the HLAAssoc-1.0 tool, which accurately infers HLA-DRB3/4/5 alleles linked to one or more HLA-DRB1 alleles. Moreover, we have demonstrated that integrating DRB3/4/5 alleles for motif deconvolution of immunopeptidome data leads to improved performance. This suggests that the implementation of HLAAssoc-1.0 in immunological studies, including MHC class II binding predictions, can serve as a valuable aid in interpreting the connections between observed HLA binding events, immune reactions, and HLA-DR allele associations. The method opens up new opportunities for studies focusing on immunotherapies, peptide vaccines, and uncovering novel HLA-DRB3/4/5 alleles that may act as risk or protective factors in autoimmune diseases. By expanding our understanding of HLA diversity and its impact on immune responses, HLAAssoc-1.0 contributes to advancing personalized medicine approaches and deepening our knowledge of human immunogenetics.

